# A novel core genome approach to enable prospective and dynamic monitoring of infectious outbreaks

**DOI:** 10.1101/421388

**Authors:** Helen van Aggelen, Raivo Kolde, Hareesh Chamarthi, Joshua Loving, Yu Fan, John T. Fallon, Weihua Huang, Guiqing Wang, Mary M. Fortunato-Habib, Juan J. Carmona, Brian D. Gross

## Abstract

Whole-genome sequencing is increasingly adopted in clinical settings to identify pathogen transmissions. Currently, such studies are performed largely retrospectively, but to be actionable they need to be carried out prospectively, in which samples are continuously added and compared to previous samples. To enable prospective pathogen comparison, genomic relatedness metrics based on single nucleotide differences must be consistent across time, efficient to compute and reliable for a large variety of samples. The choice of genomic regions to compare, *i.e.*, the *core genome*, is critical to obtain a good metric.

We propose a novel core genome method that selects conserved sequences in the reference genome by comparing its k-mer content to that of publicly available genome assemblies. The conserved-sequence genome is sample set-independent, which enables prospective pathogen monitoring. Based on clinical data sets of 3436 *S. aureus*, 1362 *K. pneumoniae* and 348 *E. faecium* samples, we show that the conserved-sequence genome disambiguates same-patient samples better than a core genome consisting of conserved genes. The conserved-sequence genome confirms outbreak samples with high accuracy: in a set of 2335 *S. aureus* samples, it correctly identifies 44 out of 45 outbreak samples, whereas the conserved gene method confirms 38 out of 45 outbreak samples.

## 1. Introduction

Genomic analysis has become an indispensable tool for identification, disambiguation, and classification of infectious agents. As the cost of whole genome sequencing has dropped to a few hundred dollars per bacterial sample,^1^ infectious disease practice is increasingly turning to genomic sequencing as a routine method to monitor infections and inform antibiotic stewardship programs.^2–6^ Healthcare-acquired infections in particular present a significant threat to patient safety, as well as a financial burden to the healthcare system.^7,8^ With whole genome sequencing analysis, it is possible to identify potential transmissions, suggest sources of transmission and provide insight into the nature and phenotypes of pathogens. The adoption of this relatively new technology calls for best practices and standards to compare microbial genomes in a manner that is consistent across time and location.

While whole genome comparison provides better resolution for pathogen disambiguation than FPGE^9^ or gene-based or multi-locus sequence typing,^10–14^ it also comes with bigger challenges.^4^ The genomic ‘distance’ between pathogens can be defined in many ways, with or without a reference genome to guide the comparison.^15^ Notable reference-free approaches include k-mer composition-based metrics and an assembly-based core genome MLST^16,17^. In this work, we will focus on alignment-based pathogen comparison, since the use of a reference genome can aid in interpretation and provide information on the location of mutations. In this paradigm, samples are aligned against a common reference genome and genomic distances are computed as the number of nucleotide differences between the consensus genomes, typically including only single nucleotide variations (SNVs).

There are several challenges in quantifying SNV distances reliably in a clinical setting. First, clinical sites typically experience a wide variety of samples and the reference genome used for the analysis – usually one per species – may be quite distant from some of the samples observed, which leads to a situation in which small differences between large numbers of SNVs need to be detected reliably. Moreover, the observed pathogens may lack some of the regions in the reference genome, in which case the comparison becomes problematic. Second, mutation rates can vary greatly across the genome^18,19^ and bias a distance metric towards hypervariable regions. Third, the distance metric needs to be sample-independent in prospective studies such that the distances between samples *a* and *b* do not change when samples *c, d*, … are added. This requirement ensures that the interpretation of sample relationships does not change over time.

The choice of core genome, *i.e*., the regions included in the genome comparison, is crucial to obtain a reliable and consistent genomic distance metric. We have developed a conserved-sequence genome that achieves these objectives, and we demonstrate this with large clinical data sets for gram-positive and gram-negative pathogens: *S. aureus*, *E. faecium* and *K. pneumoniae*. The conserved sequences that make up the core genome can be computed efficiently (without the need for multi-sequence alignment) from publicly available genome assemblies based on k-mer frequency analysis.

Our goal is to compare and evaluate this conserved-sequence method against other core genome approaches typically used for whole-genome SNV comparison. Few prior work exists on core genome approaches for applications in clinical epidemiology,^13^ so we aim to provide insight into the advantages and drawbacks of commonly used methods, including conserved-gene approaches^20^ and approaches that select genome regions with sufficient coverage in a set of samples^21–23^. We illustrate that sample-dependent core genome definitions are not suitable for prospective studies because they lead to variable SNV distances as samples are added, which complicates clinical decision-making. Conserved-gene methods, on the other hand, are suitable for prospective whole-genome comparison, as is the conserved-sequence approach we developed. We compare these approaches on several aspects: (i) their ability to separate same-patient samples from different-patient samples, (ii) their use in comparing samples across time and location, and (iii) their ability to confirm previously published outbreak samples of *S. aureus* in a neonatal ICU unit.^21^ We also detail a methodology to determine thresholds for pathogen relatedness for these core genome methods.

## 2. Results

### a. Sample-dependent core genomes yield variable genomic distances in a prospective setting

The published body of genomic epidemiology work overwhelmingly deals with retrospective studies,^21,24–27^ in which a defined set of samples is analyzed simultaneously and independently of other sample sets. As genomics is moving into the clinic as a routine methodology for infection control, a prospective mode of operation is needed to provide actionable results at any point in time that are consistent with future sample analysis. Such consistency constraints are typically not accounted for by popular methods for genomic distance calculation.

Retrospective clinical studies often compute SNV distances across an intersection of nucleotides that are unambiguously determined in all samples, an approach we will refer to as the “intersection” core genome (see Method section 4.c). This approach is practically impossible to apply in a prospective setting, because the shared regions with sufficient coverage and unambiguously determined bases decrease as the sample set grows. This implies that all distances between samples, including previously compared samples, have to be recomputed and that the resulting genomic distances decrease as samples accumulate.

To illustrate the variability of genomic distances with the intersection core genome, we simulate a prospective analysis of a *K. pneumoniae* data set^28^ by analyzing the samples incrementally (Figure 1). Each time samples are added, all sample distances have to be recomputed, which can lead to a significant computational overhead: comparing around 2000 samples in a pairwise fashion amounts to nearly 2 million core genome distances to recompute. More importantly, the SNV threshold that separates same-patient samples from different-patient samples keeps decreasing as samples are added. This highlights the problem of interpretation: the thresholds would have to be re-evaluated continuously and, even with such variable thresholds, groupings of genetically similar samples would not be guaranteed to be stable over time.

**Figure 1:**
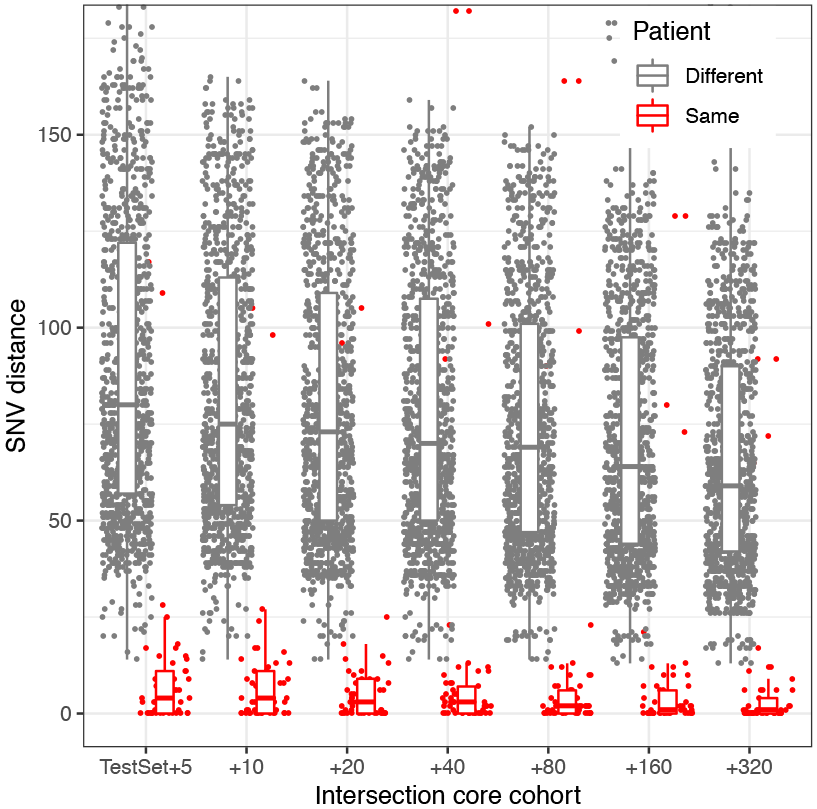
Evolution of intersection core genome SNV distances with increasing cohort size for a set of K. pneumoniae samples. Within-patient and between-patient SNV distances are shown for the same set of 49 samples as the core genome evolves when samples are added. The x axis indicates the number of samples added to the original set.

### b. Sample-independent core genome methods rely on conserved genome regions

To deliver consistent pathogen similarities and facilitate their computation in a clinical setting, the core genome must be defined from the start of the prospective study and kept fixed throughout the study. The core genome therefore needs to rely on genomic regions that are shared by the majority of strains to be compared. However, clinical settings typically see a wide variety of strains and should be able to deal with unexpected or novel strains.

A common solution is to use “housekeeping” genes, that is, genes that are highly conserved within the species, as a core genome.^29^ These genes can be identified in a sample set-independent manner by comparing gene content in publicly available finished genomes and assemblies. In this work we refer this method as “conserved-gene” core genome. A drawback of this approach is that mutation rates can vary locally, even within genes, and this can lead to an over-representation of highly variable sites within the SNV distance.

To create an SNV distance metric built on genomic sites with similar mutation rates, we developed a core genome that consists of highly conserved sequences, regardless of their gene content. We will refer to this method as the “conserved-sequence” core genome. Nucleotide conservation scores can be difficult to compute since multi-sequence alignment is computationally challenging for a large number of sequences.^30,31^ Instead of using multi-sequence alignment, we estimate the conservation of each nucleotide in the reference genome through the relative frequency of k-mers spanning the nucleotide in a set of genome assemblies (Method section 4.e). The conservation scores are computed in three steps (Figure 2). First, the reference genome and all genome assemblies are k-merized. Then, for each k-mer in the reference genome, the relative number of assemblies that contain the same k-mer is computed and associated with the k-mer’s start position *p* in the reference genome (the function ‘k-mer frequency’ in Figure 2). Finally, a conservation score for each position *p* is obtained from this function by taking a running maximum in a window [*p-k+1*, *p*] with *k* the k-mer length (the function ‘conservation score’ in Figure 2). Like the conserved-gene approach, this core genome can be composed from genome assemblies, which enables it to capture a large variety of strains while remaining sample-set independent.

**Figure 2:**
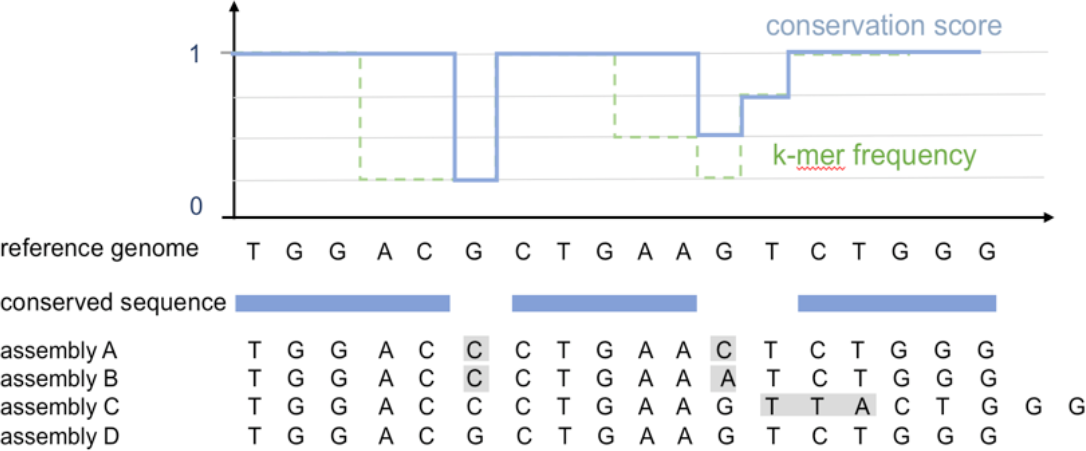
Illustration of the k-mer frequency method used to select conserved sequences in the reference genome for an example using 3-mers, a hypothetical set of 4 short genome assemblies and a frequency cutoff of 95% (in practice, longer k-mers are used to favor unique mappings to the genome). The function ‘k-mer frequency’ represents the relative frequency of the reference genome k-mers in the genome assembly set, and the function ‘conservation score’ is generated from this by taking a running maximum in an interval of 3 nucleotides.

We built conserved-gene and conserved-sequence genomes from all available RefSeq assemblies for *S. aureus*, *E. faecium* and *K. pneumoniae* (Method section 4.a) with a 95% conservation threshold for both methods. Since both of these core genome approaches rely on conserved genome regions, they largely overlap in core gene regions, but differ on a nucleotide level. While the conserved-gene core genome targets housekeeping genes that are expected to be shared by nearly all strains, the conserved-sequence core genome is not limited to gene sequences, resulting in a larger core genome (Table 1; Figure 3). The number of SNVs within the core regions is considerably smaller for the conserved-sequence than the conserved-gene cores (Figure 3), even though the latter is shorter. This reflects the success of our strategy to remove regions commonly displaying variation between strains, resulting in a focus on regions of similar mutation rate where only de novo mutations are expected.

**Figure 3:**
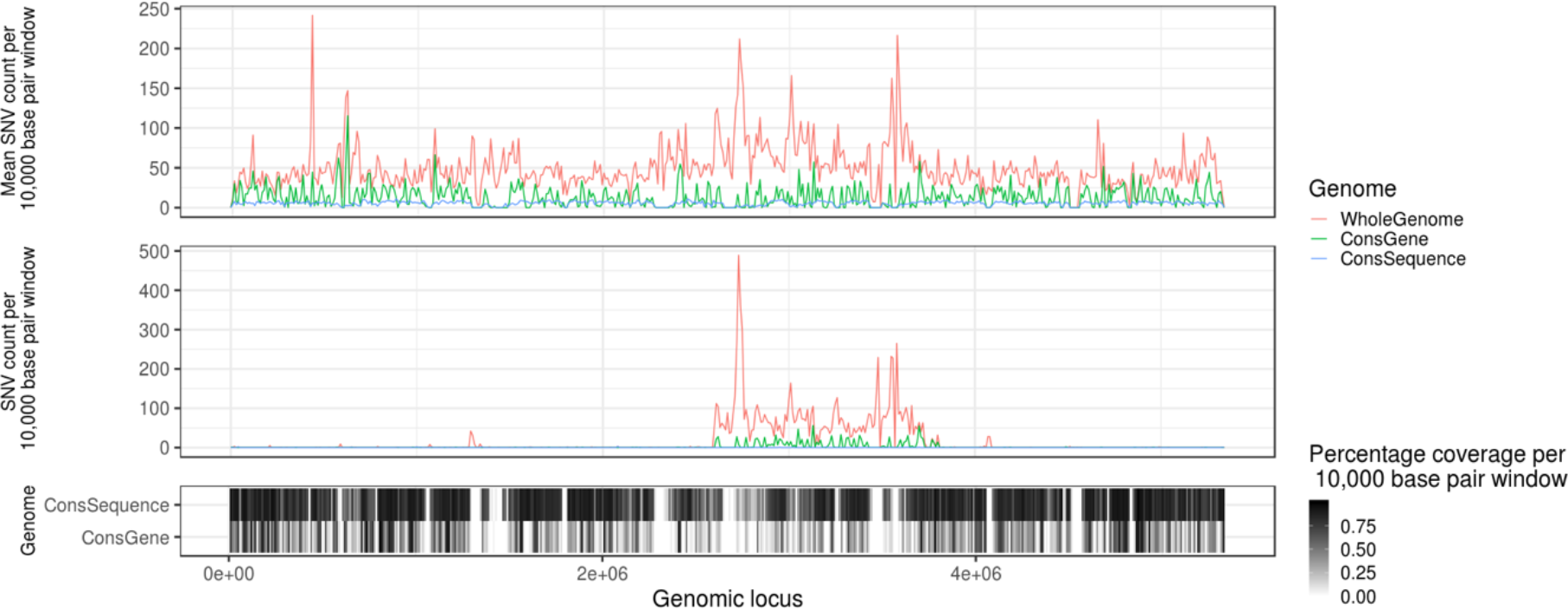
Histogram of the number of SNV counts over 10,000 base pair windows across K. pneumoniae genome, averaged across all samples (top panel) and in one random sample (middle panel). The percentage of bases covered by a core genome in the corresponding windows is shown in the bottom.

**Table 1:** Number of nucleotides and the corresponding fraction of the reference genome included in the conserved-gene and conserved-sequence core genomes.

|  | Conserved genes | Conserved sequences |
| --- | --- | --- |
| *S. aureus* | 776,385 (28%) | 1,843,071 (65%) |
| *E. faecium* | 530,487 (11%) | 994,708 (37%) |
| *K. pneumoniae* | 1,578,252 (24%) | 3,574,338 (67%) |

### c. Comparison of core genome approaches on distinction of same-patient samples

To compare the conserved-gene and conserved-sequence core genome, we evaluate their ability to separate same-patient (and same strain type) samples from different-patient samples within large clinical data sets. The data sets are not enriched for outbreak samples, so we assume that the same-patient samples largely represent same-pathogen samples. This assumption is not perfect – patients can have multiple infections of the same strain type or have a transmitted infection – but it is a useful heuristic to compare different core genome approaches on actual clinical data.

We find that the conserved-sequence core genome obtains a better separation of same-patient samples than the conserved-gene core genome, as measured by their ROC curves for large data sets for *S. aureus*, *K. pneumoniae* and *E. faecium* (Figure 4). For the large set of over three thousand *S. aureus* samples, 80% of same-patient pairs can be captured with SNV thresholds of 49, 22 and 7 for the conserved-gene, conserved-sequence and intersection method, respectively (Figure 4A). For this threshold, the conserved-gene method has the highest rate of false positives as it captures 95,464 different-patient samples, while the conserved-sequence method captures 9,605 different-patient pairs and the intersection method captures 3,292 different-patient pairs. While some of these different-patient pairs may represent actual transmissions, the disparity between these numbers suggests that at least most of the 95,464 pairs are false positives. This also illustrates how a well-chosen core genome can reduce the number of cases that need to be investigated by infection control teams.

**Figure 4:**
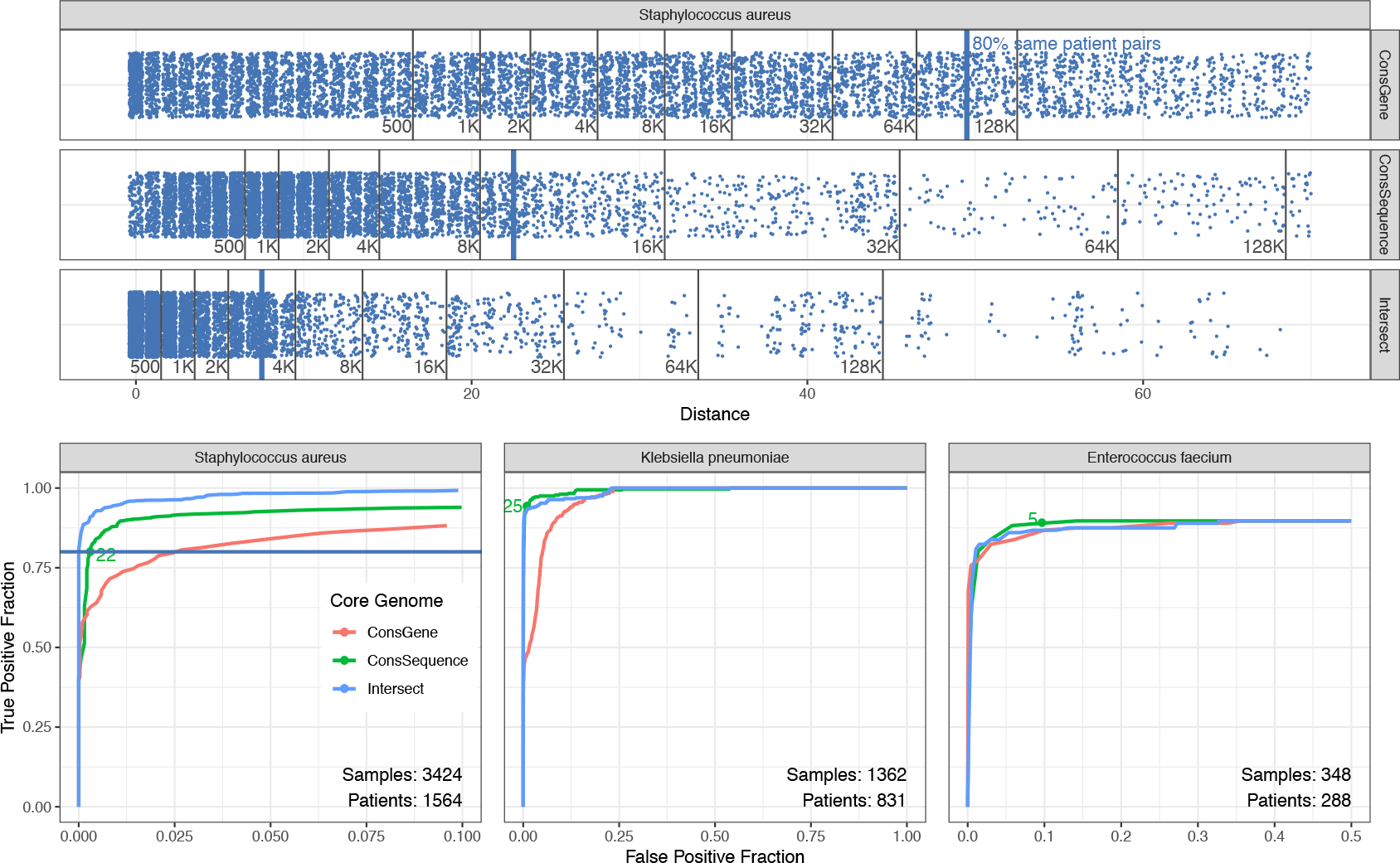
A comparison of different core genome methods to distinguish same-patient samples from different-patient samples. Panel A shows the different-patient pairs (blue dots) that fall below the SNV threshold that confirms 80% of same-patient pairs in S. aureus. The SNV thresholds correspond to 49 for the conserved-gene, 22 for conserved-sequence and 7 for intersection core genome. Panel B shows ROC curves for same-patient versus different-patient sample distinction for S. aureus (same data and 80% threshold as in panel A), K.pneumoniae and E. faecium (selected SNV threshold in green).

The 80% same-patient threshold does not necessarily represent the best true positive-false positive trade-off for each method. The ability to discriminate between same-patient and different-patient samples for different true/false positive trade-offs is summarized by the ROC curve (Figure 4B). For all species, the conserved gene method demonstrates the poorest performance. The conserved sequence method is outperformed by the intersection core genome in *S. aureus*, as illustrated for the 80% same-patient threshold, but performs better than the intersection core genome for *K. pneumoniae*. For *E. faecium*, it is difficult to detect meaningful differences since the data set is small and contains a limited number of same-patient replicates.

The ROC curves that display same-patient versus different-patient sample distinction can be used to set SNV thresholds for the detection of potential transmission clusters. The threshold should be chosen to maximize the true/false positive ratio, while prioritizing larger thresholds to enable some room for error (Figure 4B). It is important to note that these thresholds are not universal – they depend on the data used, the choice of core genome and the bioinformatic analysis pipeline used to process the samples. Given these parameters, the ROC curves can be used to set thresholds for a particular application.

### d. Benchmarking the conserved-sequence genome through verification of replicates

To evaluate whether the conserved sequence core genome approach correctly identifies same-pathogen samples – a measure of the true positive rate for transmission detection – we applied it to a set of 81 technical and biological replicates generated from three isolates for each species considered (see Method section 4.a for details). Using the SNV thresholds we described in previous section, nearly all replicates of *E. faecium*, *S. aureus*, and *K. pneumoniae* fall below the thresholds, while the inter-isolate distances lie above the thresholds (Figure 5), except for isolate pairs *a-b* and *g-h*, which are isogenic isolates. The SNV thresholds we set based on the large clinical data sets generalize well to the replicate data set. This shows that the conserved sequence approach is able to provide consistent results across a variety of strains and variation in experimental setup and sample processing, resulting in genome coverages ranging from 50x to 200x.

**Figure 5:**
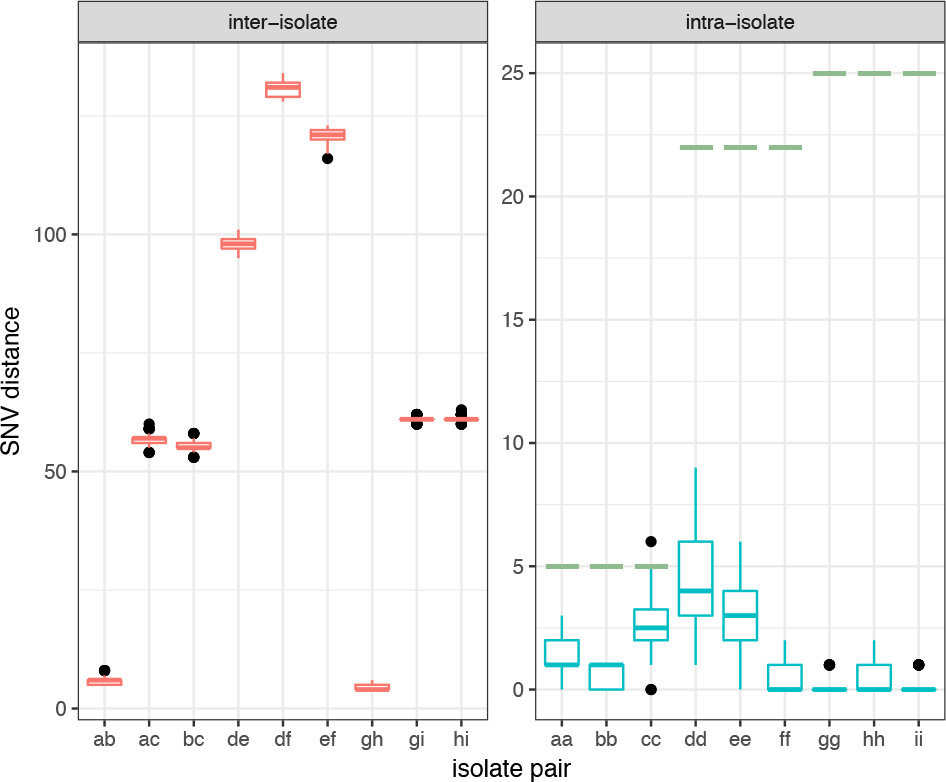
Distribution of inter-isolate (left panel) and intra-isolate SNV distances (right panel) using the conserved-sequence method between replicates for E. faecium (isolates a-c, 9 replicates each), S. aureus (isolates d-f, 9 replicates each), and K. pneumoniae (isolates g-i, 9 replicates each) represented by boxplots. The SNV thresholds for genomic relatedness for each species are indicated in green.

### e. Comparison of genomic samples across geographical locations

The conserved-sequence and conserved-gene approaches produce SNV distances that are independent of the sample set. This is not only useful to compare samples across time frames – it also makes it possible to compare samples across locations without having to anticipate the samples for all locations. Here, we test how samples compare across clinical sites with different core genome methods.

We consider *S. aureus* cohorts from multiple locations in the U.K.^32,33^ and *E. faecium* from Australia^34^ and the U.K.^35^ Since we expect to see few direct transmissions between these geographically distinct sites, the SNV distances should be clearly distinguishable from the same-patient distances for these data sets (plotted in Figure 4). We tested this for the three different core genome methods considered (Figure 6). Comparing the inter-site sample SNV distances to the quantiles obtained for the same-patient distances, we see that the *E. faecium* inter-site distances fall well below the 80% same-patient threshold for all core genome methods. For *S. aureus*, inter-site distances are closer to the 80% same-patient threshold, perhaps reflecting *S. aureus*’ presence as a community pathogen. The conserved-gene method finds many inter-site sample distances below the 80% same-patient threshold, most of which are likely false positives. The conserved-sequence and intersection core genome find no inter-site sample distances below the 80% same-patient threshold. The intersection core genome method leads to a better separation of inter-site sample distances than the conserved sequence method but is difficult to apply in a prospective manner across locations because it requires prior knowledge of all samples to define the core genome. Using a conserved sequence core genome and similar data processing pipelines across medical centers, sample comparison across sites becomes trivial. The ability to compare genomic results efficiently across medical centers and time frames enables real-time tracking of outbreaks and emerging threats.

**Figure 6:**
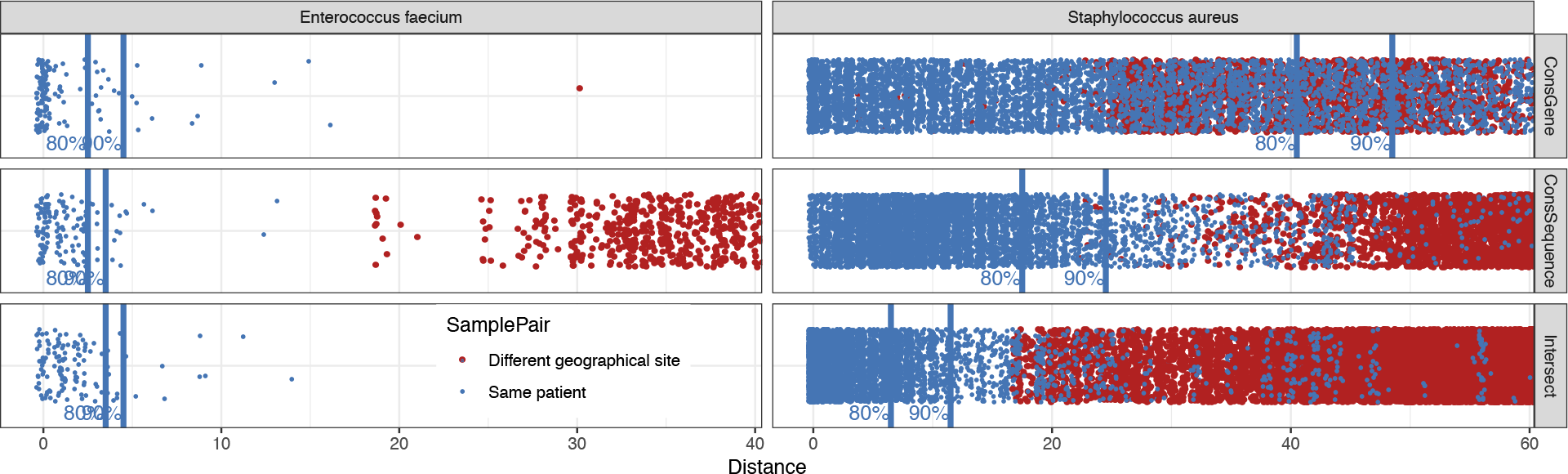
Comparison of SNV distances (in grey) between samples from geographically distinct cohorts – between S. aureus samples collected in East England and Oxford, U.K., respectively, and between E. faecium samples collected in Victoria, Australia and Cambridge, U.K. – to the quantiles obtained for same-patient distances (in red). Most of the inter-site distances fall above the 80% same-patient threshold (see Figure 4), except for the conserved gene method.

### f. Comparison of core genome approaches on a confirmed *S. aureus* outbreak

We compare the different core genome approaches’ ability to confirm and distinguish previously determined outbreak samples from a neonatal intensive care *S. aureus* study.^21^ To obtain a more realistic – and more challenging – test scenario, we add the set of 2278 East England *S. aureus* samples such that the outbreak samples present a small fraction of samples examined. As we showed in section 2.a, consistency requirements and computational considerations essentially rule out the intersection core genome for prospective application, but nonetheless represents an important benchmark. We can expect the conserved sequence and conserved gene core genome approaches to trade in some resolution relative to the intersection core genome to achieve consistency across sample sets but ideally the performance gap in a retrospective setting should be small. We defined the SNV thresholds for pathogen relatedness as the 80% same-patient threshold in the ROC curves (Figure 4).

The intersection core genome and conserved-sequence core genome find highly similar transmission clusters for the *S. aureus* outbreak samples (Figure 7). The conserved-sequence method confirms 44 out of 45 outbreak samples and finds 10 other East England samples to be similar to the outbreak; the intersection method confirms 43 and adds 9 other samples to the outbreak cluster, all 9 of which agree with those found by the conserved-sequence method. By comparison, the conserved gene method performs suboptimally, as it fails to confirm 9 of the outbreak samples. This result highlights the universal applicability of the conserved-sequence core genome: it is able to confirm the *S. aureus* outbreak samples with similar sensitivity and precision to the intersection core genome method, even though the genome regions and the SNV thresholds were defined independently of the outbreak data set. In contrast, the intersection core genome was composed from the sample data, including the outbreak samples.

**Figure 7:**
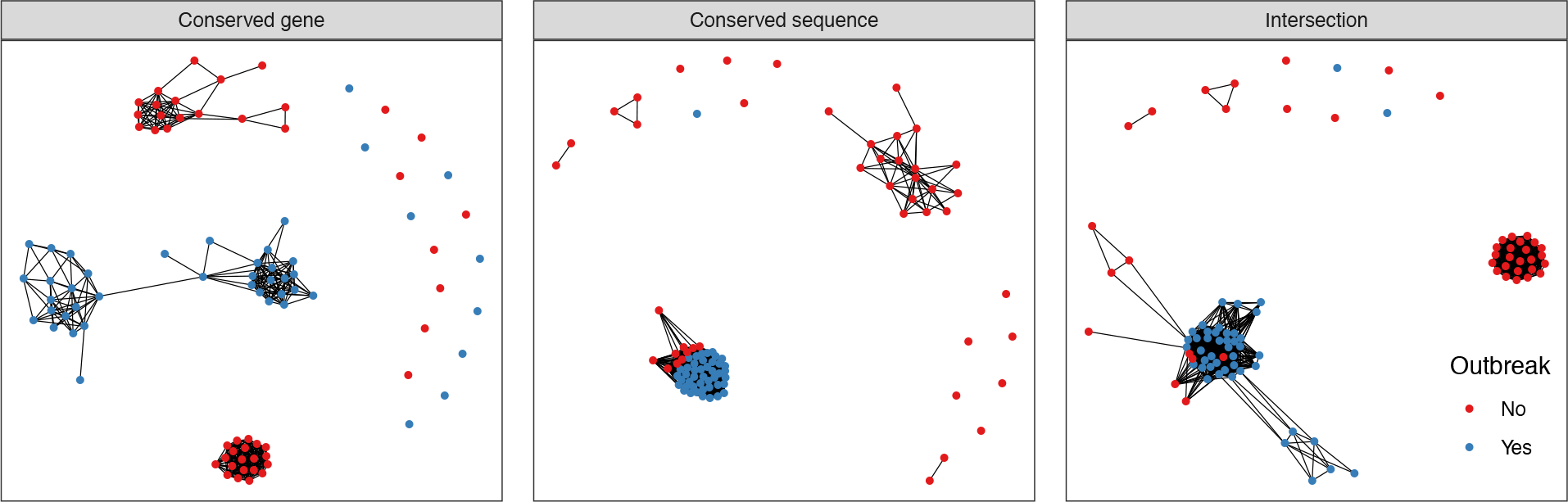
Network plot displaying single-linkage clusters of confirmed S. aureus outbreak samples (blue dots) and the 45 closest other samples (red dots) based on genomic distances computed with different core genome methods. Connected dots represent clusters of samples with SNV distances below the threshold for relatedness (i.e., the 80% same-patient threshold in Figure 4).

## 3. Discussion

Each core genome method has advantages and disadvantages and the most suitable method depends on the application. In a clinical environment, in which samples are continuously added and analyzed, stable SNV distances are needed to provide consistent insights over time and to avoid recomputing SNV distances. The commonly used intersection core genome, consisting of genome regions with high-confidence reference/variant calls in all samples, does not provide such consistency (Figure 1). Conserved-gene and conserved-sequence approaches, such as the one we introduced here, look at regions of the genome that are conserved among strains and can be applied invariably to all samples. The resulting SNV distances do not depend on the sample set and thus provide consistent genomic relationships across time and location. The ability to compare samples efficiently and dynamically within and across hospitals is crucial to identify outbreaks and emerging threats.

The conserved-sequence method we developed is suitable for dynamic clinical studies and provides superior sensitivity and specificity for pathogen relatedness over the conserved gene method. This is true for all species examined (Figure 4). Comparing *S. aureus* and *E. faecium* samples from different locations (Figure 6), the conserved-sequence method finds few small SNV distances, consistent with a low false-positive rate. The difference in accuracy between the conserved genome methods is very clear for the *S. aureus* outbreak, for which the conserved-sequence method confirms 44 out of 45 outbreak samples (Figure 7), while the conserved-gene method confirms only 36. The conserved-sequence method performs as well as the intersection core genome on this outbreak study but has the advantage that it can be applied towards prospective infectious disease epidemiology.

While the intersection core genome approach is not suitable for dynamic clinical analysis, the conserved genome approaches have the drawback that they put a stronger requirement on the consistency of variant calls across samples: inconsistent calls add to the error on the genomic distance metric in fixed core genome approaches, while they are often excluded from consideration in an intersection core genome approach because they correspond to low-confidence calls. The variability introduced by variant calling and experimental errors can be benchmarked via sample replicates (Figure 5). The ability of the core-genome SNV distances to separate same-pathogen samples from different-pathogen samples is therefore a function of the core genome method, the choice of reference genome and the bioinformatics analysis. These factors should be considered when formulating SNV thresholds for pathogen relatedness. In the absence of large same-pathogen labeled data sets, SNV thresholds can be determined via ROC curves for same-patient (same-strain-type) samples versus different-patient samples (Figure 4).

## 4. Methods

### a. Data sources

The fixed core genome approaches we evaluate build a core genome based on reference genome data, typically genome assemblies. We use all available RefSeq genome assemblies for the species considered, including 782 assemblies for *E. faecium*, 2876 for *K. pneumoniae* and 8274 for *S. aureus*.

The replicate study consists of 3 isolates for each species, obtained from Westchester Medical Center. For each isolate that is sequenced, two additional sequencing replicates, two additional library replicates, and two additional colonies from the same plate are analyzed. Two additional culture replicates (from the same colony) are sequenced one and two days after the original isolate, yielding a total of 9 replicates for each isolate. The libraries are prepared using Nextera XT DNA library preparation kits and sequenced using Illumina 2 × 150 bp reads.

All other data used are public data sets from the NCBI Sequencing Read Archive (SRA): a 57 sample set of *S. aureus* from Cambridge, U.K. (PRJEB2737),^21^ a 1158 sample set of *S. aureus* from Oxford, U.K. (PRJNA369475),^33^ a 2278 sample set of methicillin-resistant *S. aureus* (MRSA) from East England (PRJEB3174),^32^ a 1362 sample set of predominantly extended-spectrum beta-lactamase producing *K. pneumoniae* from Houston, TX (PRJNA376414),^28^ a 310 sample set of vancomycin-resistant Enterococcus (VRE) from Victoria, Australia (PRJNA433676)^34^ and a 175 sample set of VRE from Cambridge, U.K. (PRJEB12937).^35^ The VRE set from Cambridge contains longitudinal sampling and same-day replicates from a small number of patients. We remove the same-day replicates by selecting only the sample with the greatest sequencing depth for the same combination of patient ID, sequence type, sample source and sampling date. We also redefine patients as unique combinations of patient and sequence type of the pathogen in this study, since several patients had multiple distinct infections with different sequence types.

### b. Genome alignment and variant calling

All sequencing data consist of Illumina paired-end reads, which are trimmed using Trimmomatic^36^ and aligned with BWA-mem^37^ against the following reference genomes: *Staphylococcus aureus* NCTC 8325 (GenBank CP000253.1), *Enterococcus faecium* DO (GenBank CP003583.1), *Klebsiella pneumoniae* HS11286 (GenBank CP003200.1). Single nucleotide variants (SNVs) are then called with Freebayes, with a minimum base quality of 20 and subsequently filtered with vcffilter to keep SNV’s with allele frequency greater than 0.9 and a quality score greater than 5. The core genome definitions are stored in BED format, and this format is used to filter the SNV calls for the core genome regions using BCFtools’ intersection operation. SNV distances are then computed based on the consensus sequences filtered for the core genome regions. Only samples with coverage breadth > 80% and depth > 30x and fewer than 5000 ambiguous base calls (including low coverage and variant calls with < 90% allele frequencies) are included in our analysis (Supplementary Figure S1). In the *Klebsiella* dataset, we additionally remove samples for which were unable to call a multi-locus sequence type based on the variant calls.

### c. Intersection core genome

The intersection core genome approach selects the reference genome nucleotides for which all samples have sufficient read coverage and unambiguous variant/reference calls. More specifically, a position *p* in the reference genome is included in the intersection core genome if all samples considered have read coverage depth greater than 20x at this position after alignment to the reference and no sample has an ambiguous variant call in this position (*i.e*., a variant call with < 90% allele frequency).

### d. Conserved-gene core genome

The conserved-gene core genome consists of genes found in greater than 95% of all RefSeq reference assemblies considered. The genes in these assemblies are identified with Prokka.^38^ The genes are then clustered by identity and for each gene cluster, the number of assemblies containing a gene in the cluster is counted using Roary.^39^ The genes that occur in greater than 95% of assemblies are then selected as the core genome.

### e. Conserved-sequence core genome

The conserved-sequence core genome consists of conserved sequences within the reference genome. Nucleotide-based conservation scores are computed from all RefSeq assemblies for the same species via a k-mer frequency heuristic (Section 2.b). The conservation scores reflect the fraction of assemblies that contain the same nucleotide as the reference genome in a given position. To k-merize the genome assemblies, we use Jellyfish^40^, with 31 as the k-mer length. We use a 95% cutoff on the conservation score to select the core genome.

### f. Intersection core genome evolution with increasing cohort size

To visualize the intersection core genome SNV distances with increasing cohort size (Figure 1), we select all sequence type 307 samples from the Houston *K. pneumoniae* set for which the same patient was sampled at least three times (49 samples). We use the procedure described above to define the intersection core genomes, while adding increasing numbers of arbitrarily chosen samples (of any sequence type) from the same Houston cohort to the original set.

### g. ROC curves for same-patient sample distinction

To compose ROC curves for disambiguation of same-patient (positive) sample pairs, we use all datasets, except for the *S. aureus* outbreak data, combined per species. Each point on the ROC curve represents a true-positive/false-positive tradeoff with a particular SNV threshold. For *S. aureus,* we select an 80% same-patient threshold for all the core genomes to be used in subsequent analyses, yielding SNV thresholds of 49 for the conserved gene method, 22 for the conserved sequence method and 7 for the intersection core genome. For the other species, we set SNV thresholds for the conserved-sequence core genome along the bend in the ROC curve to maximize the true positive rate while minimizing the false positive rate, resulting in SNV thresholds of 25 and 5 for *K. pneumoniae* and *E. faecium*, respectively.

### h. *S. aureus* outbreak case study

For this study, we combine samples from the Cambridge outbreak^21^ with the East England MRSA study,^32^ which contains samples from Cambridge and surrounding locations. For each core genome method, the plot displays the 45 outbreak samples and the 45 other samples with the smallest distances to the outbreak samples.

## 5. Conclusion

The adoption of rapid whole-genome-based pathogen profiling for clinical epidemiology comes with a need for standards and methods that provide consistent genomic insights. Perhaps most importantly, we need methods that quantify pathogen similarity in a manner that is robust, universally applicable and suitable for dynamic analysis, in which samples are added and analyzed incrementally. This is different from a typical retrospective research setup, in which the entire sample set and the target pathogen are known and reference genomes can be chosen to be similar to the target pathogen. In a prospective clinical application, typically one reference genome is used per species and the pathogen comparison has to be robust to cover a large variety of strains within the species. As we have illustrated, prospective studies require a core genome that remains fixed throughout the duration of the study to quantify pathogen similarities consistently across time.

In this work, we introduced a conserved-sequence genome, composed by analyzing k-mer frequencies in publicly available genome assemblies, that can be applied universally to obtain consistent SNV distances across time and location. The k-mer frequency analysis used to compose the conserved-sequence genome can be applied to any set of assemblies, or even to sample data, allowing the core genome to capture the conserved sequences across a large variety of strains. Several optimizations are possible, such as using a set of assemblies stratified by strain type according to their clinical incidence.

We demonstrated that the conserved-sequence method confirms outbreak samples with similar sensitivity and specificity as the intersection core genome method, but it has the advantage that it enables prospective genomic studies. Compared with a conserved-gene approach, which is suitable for prospective studies, it finds outbreaks with much better sensitivity and specificity. This approach paves the way towards accurate, dynamic whole-genome pathogen comparison, making it possible to identify pathogen transmissions as they happen.

## Author information

H.v.A., R.K., Y.F., M.F.H., J.J.C. and B.D.G. contributed to the design of the study and editing of the manuscript. H.v.A. conceived conserved-sequence methodology, performed sample replicate analysis, created figures 2,5, and wrote the manuscript. R.K. performed statistical analyses, conceived testing methodology and created data visualizations 1,4,6,7,8. H.C. processed genomic data for public data sets. J.L. performed data analysis and plotting for figure 3. Y.F. created the conserved-gene methodology pipeline. J.T.F., W.H., G.W. executed experiments on sample replicates. All authors have read and approved the final manuscript.

## Data availability

The sequences from the replicate study have been submitted to the NCBI SRA database with the accession number PRJNA512284.

