## Supplementary figures for "A novel core genome approach to enable prospective and dynamic monitoring of infectious outbreaks"

### Supplementary material

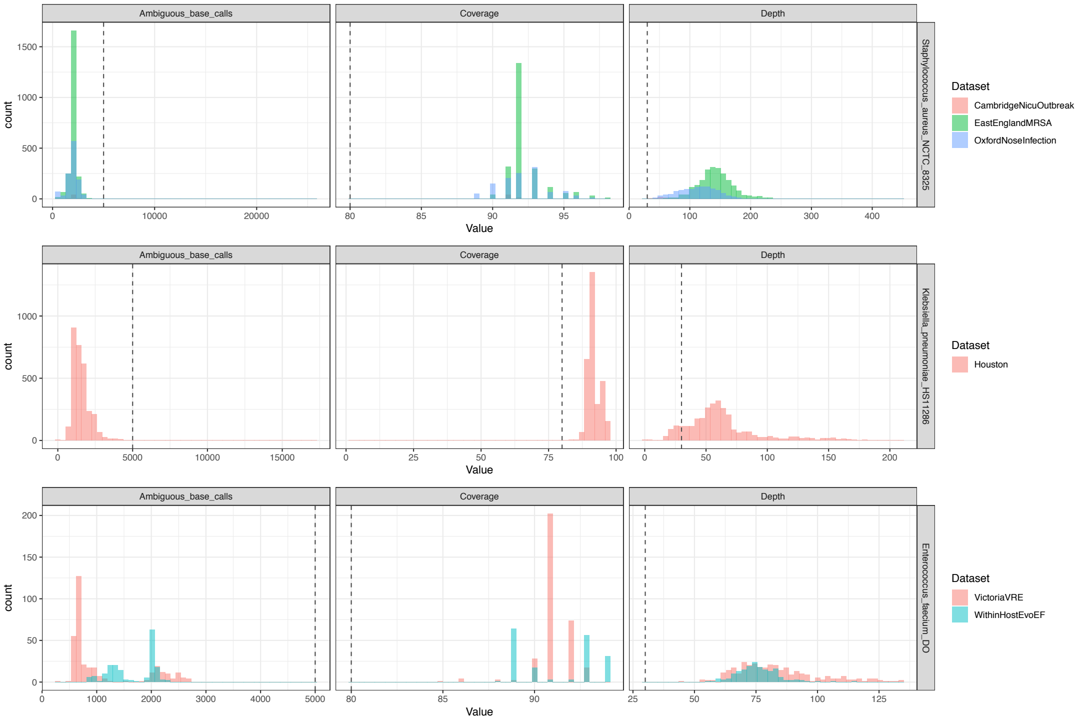


*Figure S1: Quality metrics per species (S. aureus first row, K. pneumoniae second row, E. faecium third row) for aligned data and quality thresholds used to include samples in this work.*
